## Supporting Information for "Covalent Docking and Molecular Dynamics Simulations Reveal the Specificity-Shifting Mutations Ala237Arg and Ala237Lys in TEM Beta-Lactamase"

#### Appendix S1: Choice of Docking Methodology

To measure the effects of point mutations in TEM's alanine 237 residue on the enzyme's substrate specificity, we elected to utilize Schrodinger's CovDock covalent docking workflow.[1] We defined a $\beta$-lactam opening reaction between the nucleophilic oxygen of serine 70 in TEM and the carbonyl group within the $\beta$-lactam, following previous mechanistic reports.[2] The CovDock methodology we employed is summarized in Fig S1.

As part of the CovDock workflow, we utilized Glide, which is a widely-used docking engine that employs the OPLS3 force-field[3] and a wide variety of scoring functions depending on the desired thoroughness. It uses sequential hierarchical filters to identify possible ligand binding locations within the search space, which is usually the binding site of a receptor whose shape and properties are represented on a grid.[1] Prime is a structure prediction algorithm that uses previously annotated data (when available) and *ab initio* modeling to predict the conformation of biological macromolecules. Prime's predictive accuracy is greatly increased when modeling small changes to known structures, such as point mutations in proteins.[4]

Through covalent docking, we sought to identify poses that are more likely to resemble catalytically-relevant conformations for the docked ligands, as binding modes that do not allow for the formation of a covalent bond with serine 70 are discarded after the initial scoring. Additionally, CovDock allows us to probe potential active site rearrangements in response to the binding of substrates, as structural relaxation steps are conducted after the formation of the covalent bond. This is especially important as TEM's active site is relatively occluded and might shift considerably to allow for the binding of bulkier substrates such as cefixime (Fig S3).

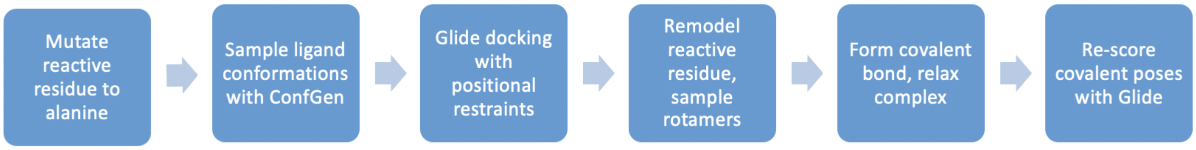


**Figure S1: Schrodinger's CovDock workflow for covalent docking, adapted from Schrodinger's documentation (release 2020-1).**

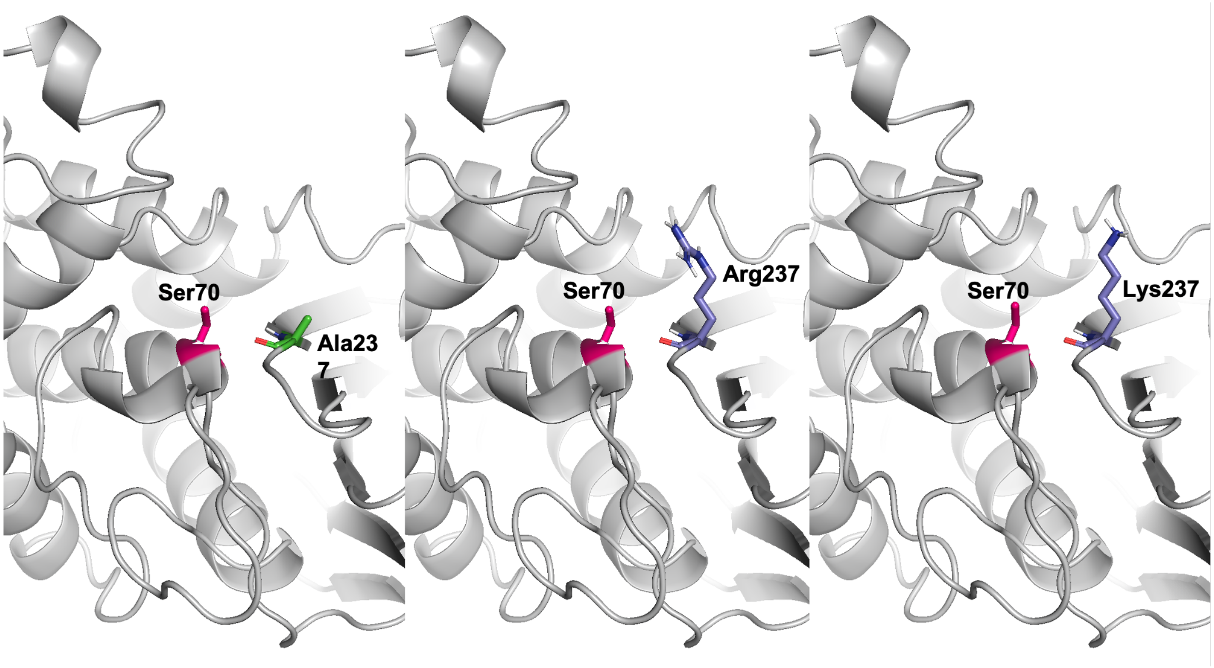


**Figure S2: Local outcome of the Ala237Arg/Lys mutations in terms of binding site occlusion. Serine 70, the catalytic residue, is represented in fuchsia.**

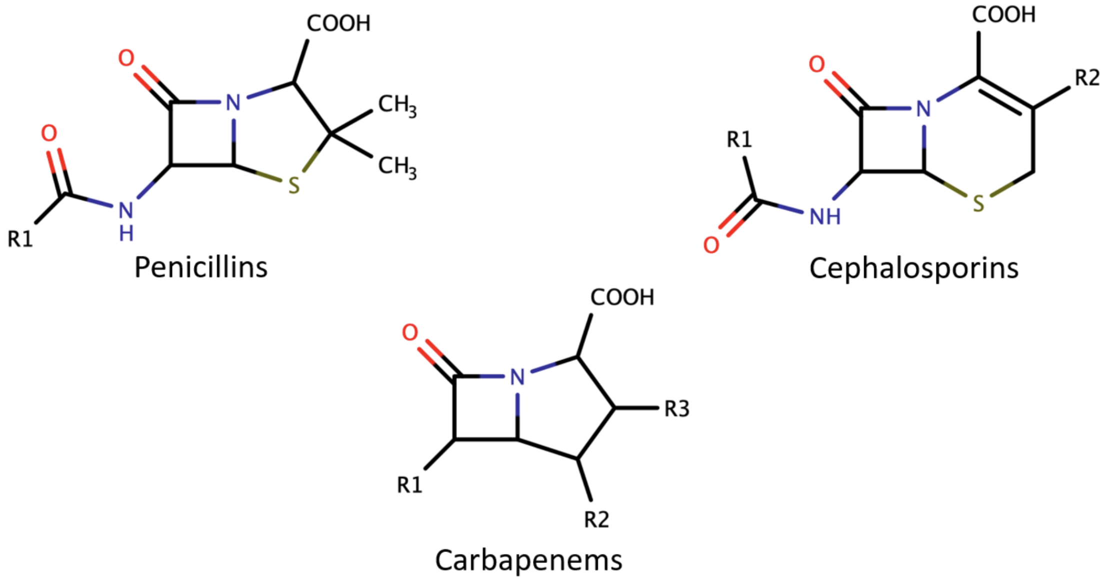


**Figure S3: Bi-dimensional representation of the scaffold shared by each of the three main drug classes tested in this study.**

**Table S2: List of all drugs used in the CovDock predictions**

| Compound | Class | PubChem CID |
|---|---|---|
| Amoxicillin | Aminopenicillin | 33613 |
| Ampicillin | Aminopenicillin | 6249 |
| Bacampicillin | Aminopenicillin | 441397 |
| Epicillin | Aminopenicillin | 71392 |
| Hetacillin | Aminopenicillin | 443387 |
| Metampicillin | Aminopenicillin | 6713928 |
| Pivampicillin | Aminopenicillin | 33478 |
| Talampicillin | Aminopenicillin | 71447 |
| Biapenem | Carbapenems | 71339 |
| Doripenem | Carbapenems | 73303 |
| Ertapenem | Carbapenems | 150610 |
| Imipenem | Carbapenems | 104838 |
| Meropenem | Carbapenems | 441130 |
| Panipenem | Carbapenems | 72015 |
| Cefacetrile | Cephalosporins1 | 91562 |
| Cefadroxil | Cephalosporins1 | 47965 |
| Cefalexin | Cephalosporins1 | 27447 |
| Cefaloglycin | Cephalosporins1 | 19150 |
| Cefalonium | Cephalosporins1 | 21743 |
| Cefaloridine | Cephalosporins1 | 5773 |
| Cefalotin | Cephalosporins1 | 6024 |
| Cefapirin | Cephalosporins1 | 30699 |
| Cefatrizine | Cephalosporins1 | 6410758 |
| Cefazaflur | Cephalosporins1 | 40240 |

| | | |
|---|---|---|
| Cefazedone | Cephalosporins1 | 71736 |
| Cefazolin | Cephalosporins1 | 33255 |
| Cefradine | Cephalosporins1 | 38103 |
| Cefroxadine | Cephalosporins1 | 5284529 |
| Ceftezole | Cephalosporins1 | 65755 |
| Cefaclor | Cephalosporins2 | 51039 |
| Cefamandole | Cephalosporins2 | 456255 |
| Cefbuperazone | Cephalosporins2 | 127527 |
| Cefminox | Cephalosporins2 | 71141 |
| Cefonicid | Cephalosporins2 | 43594 |
| Ceforanide | Cephalosporins2 | 43507 |
| Cefotetan | Cephalosporins2 | 53025 |
| Cefoxitin | Cephalosporins2 | 441199 |
| Cefprozil | Cephalosporins2 | 5281006 |
| Cefuroxime | Cephalosporins2 | 5479529 |
| Cefuroxime Axetil | Cephalosporins2 | 6321416 |
| Cefuzonam | Cephalosporins2 | 6336505 |
| Loracarbef | Cephalosporins2 | 5284585 |
| Cefcapene | Cephalosporins3 | 6436055 |
| Cefdaloxime | Cephalosporins3 | 9571072 |
| Cefdinir | Cephalosporins3 | 6915944 |
| Cefetamet | Cephalosporins3 | 5487888 |
| Cefixime | Cephalosporins3 | 5362065 |
| Cefmenoxime | Cephalosporins3 | 9570757 |
| Cefodizime | Cephalosporins3 | 5361871 |
| Cefoperazone | Cephalosporins3 | 44187 |
| Cefotaxime | Cephalosporins3 | 5742673 |

| | | |
|---|---|---|
| Cefotiam | Cephalosporins3 | 43708 |
| Cefpimizole | Cephalosporins3 | 68597 |
| Cefsulodin | Cephalosporins3 | 656575 |
| Ceftazidime | Cephalosporins3 | 5481173 |
| Cefteram | Cephalosporins3 | 6537431 |
| Ceftibuten | Cephalosporins3 | 5282242 |
| Ceftiolene | Cephalosporins3 | 6537430 |
| Flomoxef | Cephalosporins3 | 65864 |
| Latamoxef | Cephalosporins3 | 47499 |
| Cefepime | Cephalosporins4 | 5479537 |
| Cefozopran | Cephalosporins4 | 9571080 |
| Cefpirome | Cephalosporins4 | 5479539 |
| Cefquinome | Cephalosporins4 | 5464355 |
| Ceftaroline Fosamil | Cephalosporins5 | 9852981 |
| Ceftobiprole | Cephalosporins5 | 135413542 |
| Ceftolozane | Cephalosporins5 | 53234134 |
| Aztreonam | Monobactams | 5742832 |
| Carumonam | Monobactams | 6540466 |
| NocardicinA | Monobactams | 6419429 |
| Tigemonam | Monobactams | 9576769 |
| Azidocillin | Penicillins1 | 15574941 |
| Benzathine Benzylpenicillin | Penicillins1 | 15232 |
| Benzylpenicillin | Penicillins1 | 5904 |
| Clometocillin | Penicillins1 | 71807 |
| Penamecillin | Penicillins1 | 10250769 |
| Pheneticillin | Penicillins1 | 272833 |
| Phenoxymethyl Penicillin | Penicillins1 | 6869 |

| | | |
|---|---|---|
| Procaine Benzylpenicillin | Penicillins1 | 5903 |
| Propicillin | Penicillins1 | 92879 |
| Cloxacillin | Penicillins2 | 6098 |
| Methicillin | Penicillins2 | 6087 |
| Nafcillin | Penicillins2 | 8982 |
| Oxacillin | Penicillins2 | 6196 |
| Azlocillin | Penicillins4 | 6479523 |
| Carbenicillin | Penicillins4 | 20824 |
| Carindacillin | Penicillins4 | 93184 |
| Mezlocillin | Penicillins4 | 656511 |
| Piperacillin | Penicillins4 | 43672 |
| Temocillin | Penicillins4 | 171758 |
| Ticarcillin | Penicillins4 | 36921 |

**Table S1: Sequences for the mutagenesis primers used to generate the 20 TEM Ala237X constructs.**

| Mutation | Mut_F_Primer |
|---|---|
| Ala237Arg | ACCCACGCTCACCGCGTCCAGATTTATCAGCAATAAAC |
| Ala237Asn | GACCCACGCTCACCGTTTCCAGATTTATCAGCAATAAACCA |
| Ala237Asp | GACCCACGCTCACCGTCTCCAGATTTATCAG |
| Ala237Cys | GACCCACGCTCACCGCATCCAGATTTATCAGCAATAAACCA |
| Ala237Gln | GAGACCCACGCTCACCCTGTCCAGATTTATCAGCAATAAACCA |
| Ala237Glu | GAGACCCACGCTCACCCTCTCCAGATTTATCAGC |
| Ala237Gly | GACCCACGCTCACCGCCTCCAGATTTATCAG |
| Ala237His | GACCCACGCTCACCGTGTCCAGATTTATCAGCAATAAACCA |
| Ala237Ile | GACCCACGCTCACCGATTCCAGATTTATCAGCAATAAACCA |
| Ala237Leu | CGAGACCCACGCTCACCTAGTCCAGATTTATCAGCAATAAACCAGC |
| Ala237Lys | CGAGACCCACGCTCACCCTTTCCAGATTTATCAGCAATAAACCAGC |
| Ala237Met | CGAGACCCACGCTCACCCATTCCAGATTTATCAGCAATAAACCAGC |
| Ala237Phe | GACCCACGCTCACCGAATCCAGATTTATCAGCAATAAACCA |
| Ala237Pro | CCACGCTCACCGGGTCCAGATTTATCAGCAATAAAC |
| Ala237Ser | GACCCACGCTCACCGCTTCCAGATTTATCAGCAATAAACCA |
| Ala237Thr | CCACGCTCACCGGTTCCAGATTTATCAGCAATAAAC |
| Ala237Trp | GAGACCCACGCTCACCCCATCCAGATTTATCAGCAATAAACCA |
| Ala237Tyr | CGAGACCCACGCTCACCATATCCAGATTTATCAGCAATAAACCAGC |
| Ala237Val | GACCCACGCTCACCGACTCCAGATTTATCAG |

| Mutation | Mut_R_Primer |
|---|---|
| Ala237Arg | GTTTATTGCTGATAAATCTGGACGCGGTGAGCGTGGGT |
| Ala237Asn | TGGTTTATTGCTGATAAATCTGGAAACGGTGAGCGTGGGTC |
| Ala237Asp | CTGATAAATCTGGAGACGGTGAGCGTGGGTC |
| Ala237Cys | TGGTTTATTGCTGATAAATCTGGATGCGGTGAGCGTGGGTC |
| Ala237Gln | TGGTTTATTGCTGATAAATCTGGACAGGGTGAGCGTGGGTCTC |
| Ala237Glu | GCTGATAAATCTGGAGAGGGTGAGCGTGGGTCTC |
| Ala237Gly | CTGATAAATCTGGAGGCGGTGAGCGTGGGTC |
| Ala237His | TGGTTTATTGCTGATAAATCTGGACACGGTGAGCGTGGGTC |
| Ala237Ile | TGGTTTATTGCTGATAAATCTGGAATCGGTGAGCGTGGGTC |
| Ala237Leu | GCTGGTTTATTGCTGATAAATCTGGACTAGGTGAGCGTGGGTCTCG |
| Ala237Lys | GCTGGTTTATTGCTGATAAATCTGGAAAGGGTGAGCGTGGGTCTCG |
| Ala237Met | GCTGGTTTATTGCTGATAAATCTGGAATGGGTGAGCGTGGGTCTCG |
| Ala237Phe | TGGTTTATTGCTGATAAATCTGGATTCGGTGAGCGTGGGTC |
| Ala237Pro | GTTTATTGCTGATAAATCTGGACCCGGTGAGCGTGG |
| Ala237Ser | TGGTTTATTGCTGATAAATCTGGAAGCGGTGAGCGTGGGTC |
| Ala237Thr | GTTTATTGCTGATAAATCTGGAACCGGTGAGCGTGG |
| Ala237Trp | TGGTTTATTGCTGATAAATCTGGATGGGGTGAGCGTGGGTCTC |
| Ala237Tyr | GCTGGTTTATTGCTGATAAATCTGGATATGGTGAGCGTGGGTCTCG |
| Ala237Val | CTGATAAATCTGGAGTCGGTGAGCGTGGGTC |

**Table S3: CovDock scores (in kcal/mol) for the three compounds investigated in detail in this study against TEM-1 and TEM Ala237Arg/Lys.**

| CovDock Scores (kcal/mol) | TEM-1 | TEM-Ala237Arg | TEM-Ala237Lys |
|---|---|---|---|
| **Ampicillin** | -6.308 | -3.874 | -4.479 |
| **Cefixime** | -3.559 | -5.403 | -5.52 |
| **Ceftibuten** | -4.99 | -6.465 | -6.681 |
| **Carumonam** | -3.256 | -4.595 | -3.817 |

**Table S4: Relative docking scores and fitness values for all tested mutants upon treatment with wither ampicillin or cefixime.** Mutants where no docking score was calculated due to extremely unfavorable poses had their scores set as ”NAN”.

| Mutation | Rel. Docking Score | Rel. Fitness | Drug |
|---|---|---|---|
| A237C | 1.031547 | 0.5 | Ampicillin |
| A237D | 0.771084 | 0.25 | Ampicillin |
| A237E | 0.950698 | 0.25 | Ampicillin |
| A237F | NAN | 0.125 | Ampicillin |
| A237G | 0.934686 | 1 | Ampicillin |
| A237H | NAN | 0.125 | Ampicillin |
| A237I | 0.907261 | 0.25 | Ampicillin |
| A237K | 0.710051 | 0.125 | Ampicillin |
| A237L | 0.980659 | 1 | Ampicillin |
| A237M | 0.882091 | 0.5 | Ampicillin |
| A237N | 0.954661 | 0.5 | Ampicillin |
| A237P | 0.462904 | 0.0625 | Ampicillin |
| A237Q | 0.911699 | 0.25 | Ampicillin |
| A237R | 0.614141 | 0.125 | Ampicillin |
| A237S | 1.004756 | 1 | Ampicillin |
| A237T | 0.987952 | 2 | Ampicillin |
| A237V | 1.012841 | 0.5 | Ampicillin |
| A237W | -0.05696 | 0.5 | Ampicillin |
| A237Y | NAN | 0.25 | Ampicillin |
| A237C | 1.198388 | 1 | Cefixime |
| A237D | 0.913865 | 1 | Cefixime |
| A237E | 1.078911 | 1 | Cefixime |
| A237F | 0.894971 | 1 | Cefixime |
| A237G | 1.044735 | 1 | Cefixime |
| A237H | 1.087802 | 1 | Cefixime |
| A237I | 1.155599 | 1 | Cefixime |
| A237K | 1.533759 | 2 | Cefixime |
| A237L | 1.298138 | 1 | Cefixime |
| A237M | 1.01923 | 1 | Cefixime |
| A237N | 1.36788 | 1 | Cefixime |
| A237P | -0.05696 | 1 | Cefixime |
| A237Q | 1.351487 | 1 | Cefixime |
| A237R | 1.50125 | 4 | Cefixime |
| A237S | 1.103362 | 1 | Cefixime |
| A237T | 1.234787 | 2 | Cefixime |
| A237V | 1.562934 | 1 | Cefixime |
| A237W | NAN | 1 | Cefixime |
| A237Y | 1.061684 | 1 | Cefixime |

**Supporting Folder 1. Input coordinates (PDB format) for structures used in this study along with README file with further details.**

### Graphical TOC Entry
